# Network topology differentially shapes ecological processes across scales in experimental metacommunities

**DOI:** 10.64898/2026.04.17.719207

**Authors:** Paulina A. Arancibia

## Abstract

The spatial configuration of habitat patches is a key driver of metacommunity dynamics, yet the role of network topology remains poorly understood. In this study, I experimentally tested how different aspects of network structure influence metacommunity processes operating at different spatial scales. Using protist microcosms, I assembled metacommunities with patches connected as random or scale-free networks, and quantified occupancy, biomass, and extinction dynamics in relation to local (patch degree) and global (closeness centrality) metrics of connectivity.

Scale-free metacommunities supported higher occupancy and biomass than random networks. At local scales, biomass declined with increasing patch degree, suggesting that reduced connectivity may enhance productivity, likely by limiting negative interactions. In contrast, extinction dynamics were not related to degree but strongly associated with patch centrality, with network topology modulating the relationship. These results reveal a decoupling between ecological processes, showing that different components of network structure can regulate dynamics at different spatial scales.

## Introduction

Most natural habitats are currently under increasing anthropogenic influence, making the ability to predict community responses an urgent challenge. Habitat loss and fragmentation in particular, can alter the spatial configuration of communities and currently represent some of the major threats for biodiversity (Haddad *et al*. 2015; Tilman *et al*. 1994). Understanding the dynamics of species under different spatial scenarios is essential to anticipate how ecological communities will respond to global change.

Metacommunity Theory (MT) provides a framework to address these questions by defining (meta)communities as networks of local assemblages connected by the dispersal of multiple potentially interacting species (Hanski & Gilpin 1991; Holyoak *et al*. 2005; Leibold *et al*. 2004). Within this framework, dynamics emerge from the interplay between dispersal, species interactions, and environmental variability. Dispersal can greatly affect metacommunity dynamics because it modulates species coexistence (Amarasekare 2003; Massol *et al*. 2017), often generating a variety of outcomes ranging from enhanced species persistence (Holyoak & Lawler 1996; Warren 1996) to negative effects on species richness (Cadotte & Fukami 2005) depending on environmental conditions.

One of the main challenges in metacommunity ecology is understanding how spatial structure shapes metacommunity dynamics. Network structure determines the pattern of connectivity between patches, and as such, it determines how individuals, species, and interactions are distributed in space. Patch size and habitat heterogeneity are key components of spatial structure and have been widely recognized as important determinants of biodiversity, particularly after the development of the theory of island biogeography (MacArthur & Wilson 1967). Since then, studies have included different aspects of spatial structure and their results have highlighted the importance of patch connectivity for the maintenance of diversity in spatially connected habitats (Gilbert *et al*. 1998; Gonzalez *et al*. 1998; Tews *et al*. 2004). Habitat connectivity, and consequently isolation, can have different magnitude effects in different taxa (Driscoll & Lindenmayer 2009; Frisch *et al*. 2012; Santangelo *et al*. 2021), and at distinct spatial scales (Moritz *et al*. 2013), making their importance in metacommunity studies very clear. For example, we know that at the local level, isolation can reduce alpha diversity (Gonzalez *et al*. 1998; Vanschoenwinkel *et al*. 2007), but its impacts on regional diversity in different communities, and under scenarios with different levels of complexity, become increasingly hard to predict.

Although spatial structure has been shown to alter the strength and timing of species interactions (Gonzalez *et al*. 2011) and consequently of species sorting, the role of network topology –the configuration of inter-patch connections within a network– has received little attention in the context of metacommunity dynamics. Insights from single-species metapopulations suggest that topology can strongly influence persistence: Gilarranz & Bascompte (2012) theoretically demonstrated that its effects on metapopulation persistence can be modulated by factors governing local extinction and colonization. In metacommunities, species interactions such as competition or predation directly shape these processes by increasing local extinction risk.

At the same time, topology is likely to influence not only the magnitude but also the spatial distribution of interactions. Because interactions are not homogeneous across patches, network structure can generate transient heterogeneities that can alter population dynamics and, in some cases, promote stability particularly in predator-prey dynamics (Holt 2002). The pattern of connectivity of patches or the network topology of a metacommunity, can have important repercussions in its dynamics (Holland & Hastings 2008), altering species richness and biomass (Chisholm *et al*. 2011; Economo & Keitt 2007) and overall stability (Arancibia 2024).

Ecological processes in metacommunities operate across spatial scales, suggesting that different aspects of network topology may regulate different processes. Different network metrics capture connectivity at distinct spatial scales. Local connectivity measured as patch degree (number of direct connections) reflects immediate dispersal inputs. In contrast, global measures such as closeness centrality, which capture the position of a patch within a network and its accessibility through indirect pathways. Although studies suggest that network topology can influence metapopulation persistence (Arancibia & Morin 2022; Gilarranz & Bascompte 2012), it remains unclear how these different aspects of network structure can regulate different ecological processes in metacommunities.

Ecological dynamics in metacommunities operate across different levels of organization. For example, measures of biomass represent community functioning and integrate processes such as interactions, resource use, growth, etc., at local scales. In contrast, a patch probability of extinction represents community persistence capturing the balance between local extinction and colonization dynamics. Occupancy dynamics, on the other hand, integrate these aspects by reflecting dispersal efficiency and the outcomes of environmental filtering.

Given the complexity and scale of metacommunities, studies aimed at testing how network topology influences multiple ecological processes remain limited. Most studies in this topic rely heavily on mathematical models, which have been extremely useful to explore mechanistic insights into metacommunity dynamics, but in most cases lack the comparison to observational or experimental data (Logue *et al*. 2011). Models have the advantage of allowing us to represent scenarios with large numbers of patches and control many variables; however, many models must introduce simplifying assumptions to balance realism and computational tractability. Producing empirical data at this scale can nonetheless be impractical and expensive, and most studies that can provide it, lack a temporal component (Brown *et al*. 2017), artificially force dispersal, or use bulk dispersal patterns that eliminate intraspecific differences in dispersal (Grainger & Gilbert 2016; Guzman *et al*. 2019; Sullivan *et al*. 2020). These simplifications greatly reduce the opportunity to draw generalizations to other systems. Controlled experiments, however, allow the analysis of certain assumptions and the testing of others under more generalized scenarios.

This study examines how network topology affects metacommunity dynamics at both regional and local scales. Because ecological processes such as occupancy, production, and extinction operate at different spatial scales, I hypothesize that they respond to different components of network topology. Specifically, I predict that local connectivity (patch degree) will primarily influence local processes such as biomass, while global connectivity (closeness centrality) will have a stronger influence on extinction dynamics. Since network topology determines the distribution of patches and their pattern of connectivity, I expect it to modulate these relationships, producing context-dependent outcomes.

## Methods

### Experimental setup

To represent two contrasting and biologically plausible patterns of spatial distribution of habitat patches in metacommunities, I designed experimental layouts with two types of network topology: Erdös-Rényi (from here on “random”) and scale-free. I used the R package ‘igraph’ (v. 1.2.4.1) to create two different —but comparable— realizations of each topology with 24 patches each (Fig. 1). In order to characterize the patches and their location in the network I used two metrics that have been previously used to describe node position/importance within a metacommunity network (Borthagaray *et al*. 2015a, b; Estrada & Bodin 2008) degree and closeness centrality. Degree corresponds to the number of links between a focal node and other nodes in the network. Closeness centrality is measured as the reciprocal of the average shortest path length between a focal node and all the other nodes in the network. Therefore, higher values indicate nodes that are closer to others or “more central”.

**Fig. 1.**
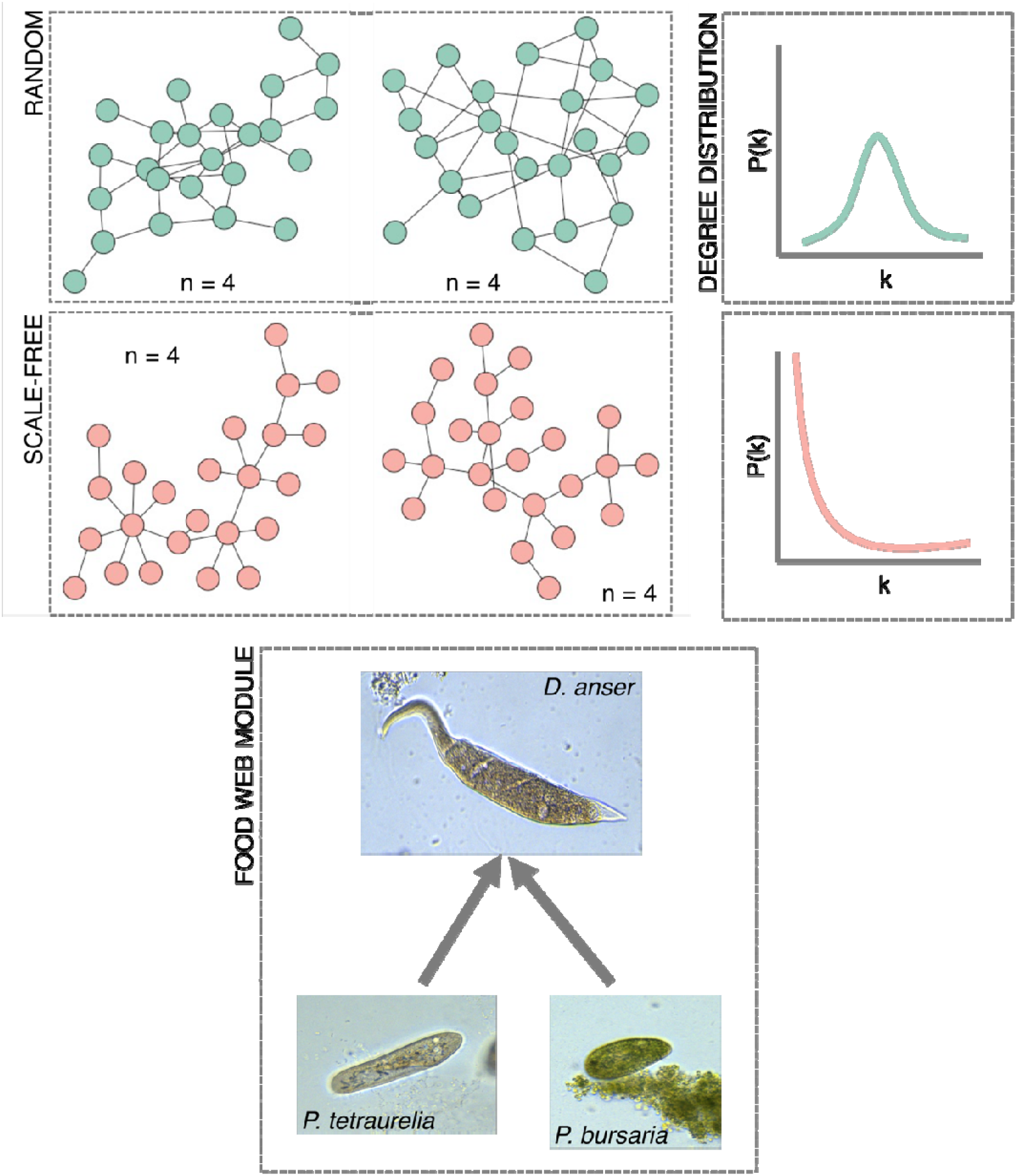
Experimental design. The top panels show the layouts of random and scale-free networks used in the experiment, and their idealized degree distribution on the right. Each network realization (i.e. layout) was replicated four times in the experiment. Bottom panel describes the food web module used in the experiment. The ciliate *Dileptus anser* preys on both *Paramecium tetraurelia* and *P. bursaria*.

The experimental metacommunities were created using 24 well-plates filled with protist media (0.37 g Carolina Protist pellet, 0.07 g Herptivite in 1.4 L of well water) previously inoculated with four bacterial species (as a food source): *Proteus vulgaris, Serratia marcesens, Bacillus subtilis*, and *Bacillus cereus*. I tracked the dynamics of three species of ciliates: *Paramecium tetraurelia, Paramecium bursaria* and *Dileptus anser*. In this food-web module, the two *Paramecium* species compete for food resources (bacteria), although the mixotroph *P. bursaria* can obtain additional energy from its mutualistic endosymbiont (*Chlorella* sp.). *Dileptus anser* is an obligate predator that feeds on both *Paramecium* species. The two prey species (*Paramecium*) were separately inoculated at random in 30% of the patches on day one of the experiment, while the predator *Dileptus* was introduced on day five to allow the prey populations to establish. Patches were connected using glass capillary tubes of similar length, filled with protist media. This allowed me to capture the differences between the natural migration patterns of the three species instead of artificially forcing them by pipetting (see Kerr *et al*. 2006 for an example). The microcosms were placed in an incubator at 24°C with a 12:12h photoperiod and were allowed to develop for 80 days (∼80 protist generations). Since all three species are morphologically distinct, I was able to evaluate their abundance in each patch by counting directly using a stereomicroscope, three times a week.

### Descriptors of metacommunity dynamics

I evaluated changes in metacommunity dynamics by estimating community biomass. For this, I used photographs of 10-35 individuals of each species and estimated their surface area using ImageJ (Schneider *et al*. 2012). I calculated biomass using surface area estimations, the value of the density of water, and the abundance counts per patch. This approach has been used previously as a proxy for biomass in protist systems where direct measurements are impractical. Metacommunity occupancy was calculated as the percentage of patches occupied at a single time point. To get insights into the probability of extinction in any given patch I calculated incidence as the percent of time steps a patch was occupied, and I interpreted the quantity (1-incidence) to represent probability of extinction (Gilarranz & Bascompte 2012).

### Statistical analyses

To evaluate differences between types of networks in occupancy and biomass, I used profile analysis (R package ‘profileR’ v. 0.3.5). This technique detects differences in parallelism and equality of response levels (Desjardins & Bulut 2015). The relationship between local response variables (biomass and patch extinction probability) and network metrics (degree and closeness centrality) was analyzed using generalized mixed models (GLMMs) with sampling day included as repeated measures, and plate and network replicate as random effects.

## Results

The three species coexisted in the metacommunity for the first 30 days of the experiment (∼30 protist generations); therefore, all the analyses were carried out considering only that period. Occupancy in both types of metacommunities increased quickly. However, after a few generations, occupancy in random metacommunities started dropping faster than in scale free. The analysis confirmed that scale-free networks showed overall higher occupancy than random (*F*_1, 14_ =16.95, *p* = 0.0015; Fig. 2). Metacommunity-level biomass data were first reported in Arancibia (2024), and showed a very similar temporal pattern to occupancy, where scale-free metacommunities harbored more organisms than random networks (Fig. S1).

**Figure 2.**
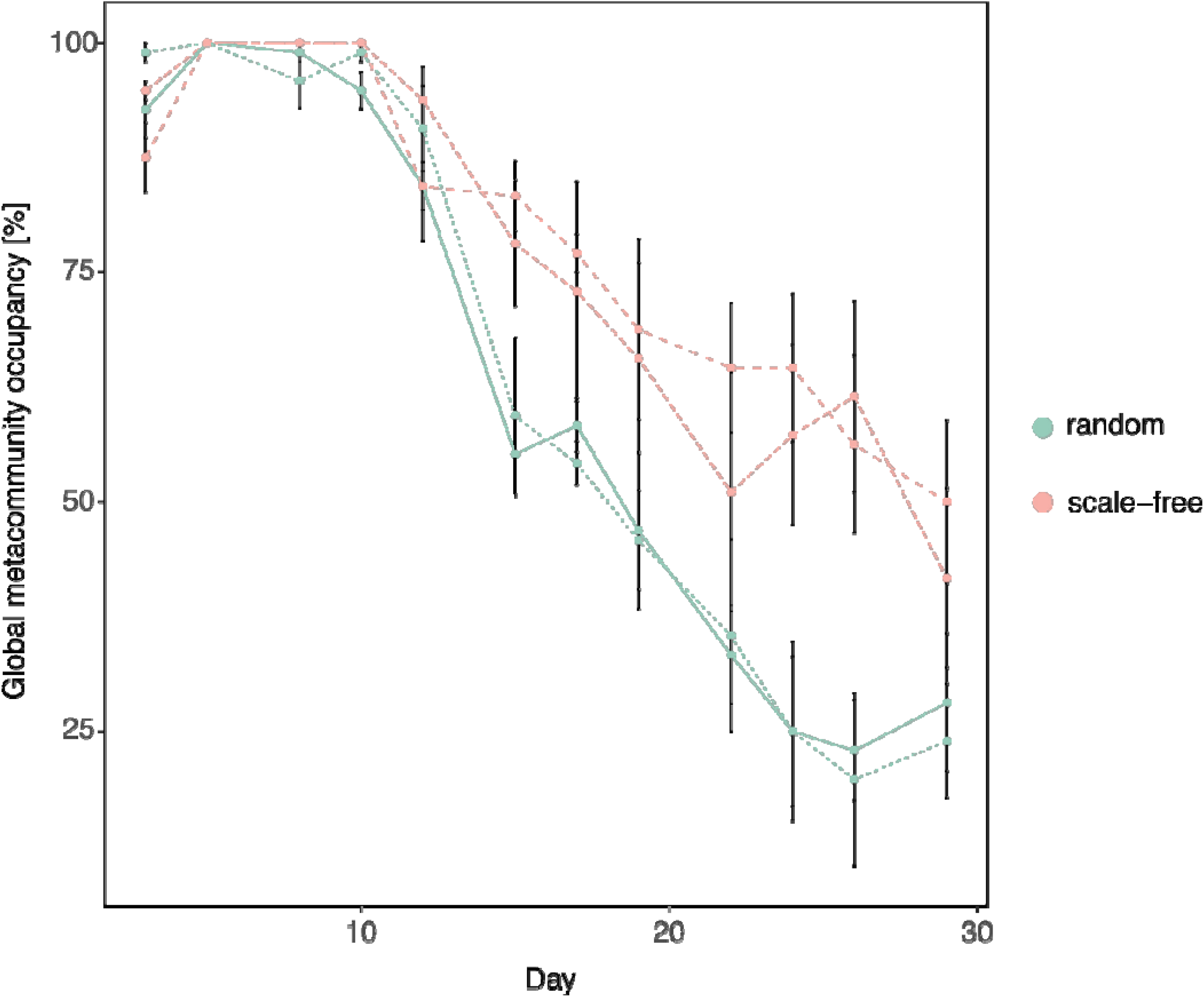
Metacommunity occupancy (% of occupied patches) over time in experimental metacommunities connected as random (blue) and scale-free (orange) networks 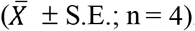. Same color lines correspond to network-level replicates.

The predator *D. anser* was the first species to go globally extinct. *Paramecium tetraurelia* had both higher occupancy (*F*_1, 14_= 15.21, *p* = 0.0016; Fig. 3A) and biomass in scale-free networks (*F*_1, 14_ = 17.59, *p* < 0.0009; Fig. 3D). *Paramecium bursaria* and *D. anser*, did not show differences between network types in occupancy patterns (Fig. 3B-C), but reached higher biomass in scale-free networks (*F*_1, 14_= 4.745, *p* = 0.047 and *F*_1, 14_ = 5.685, *p* = 0.0318 respectively; Fig. 3E-F).

**Figure 3.**
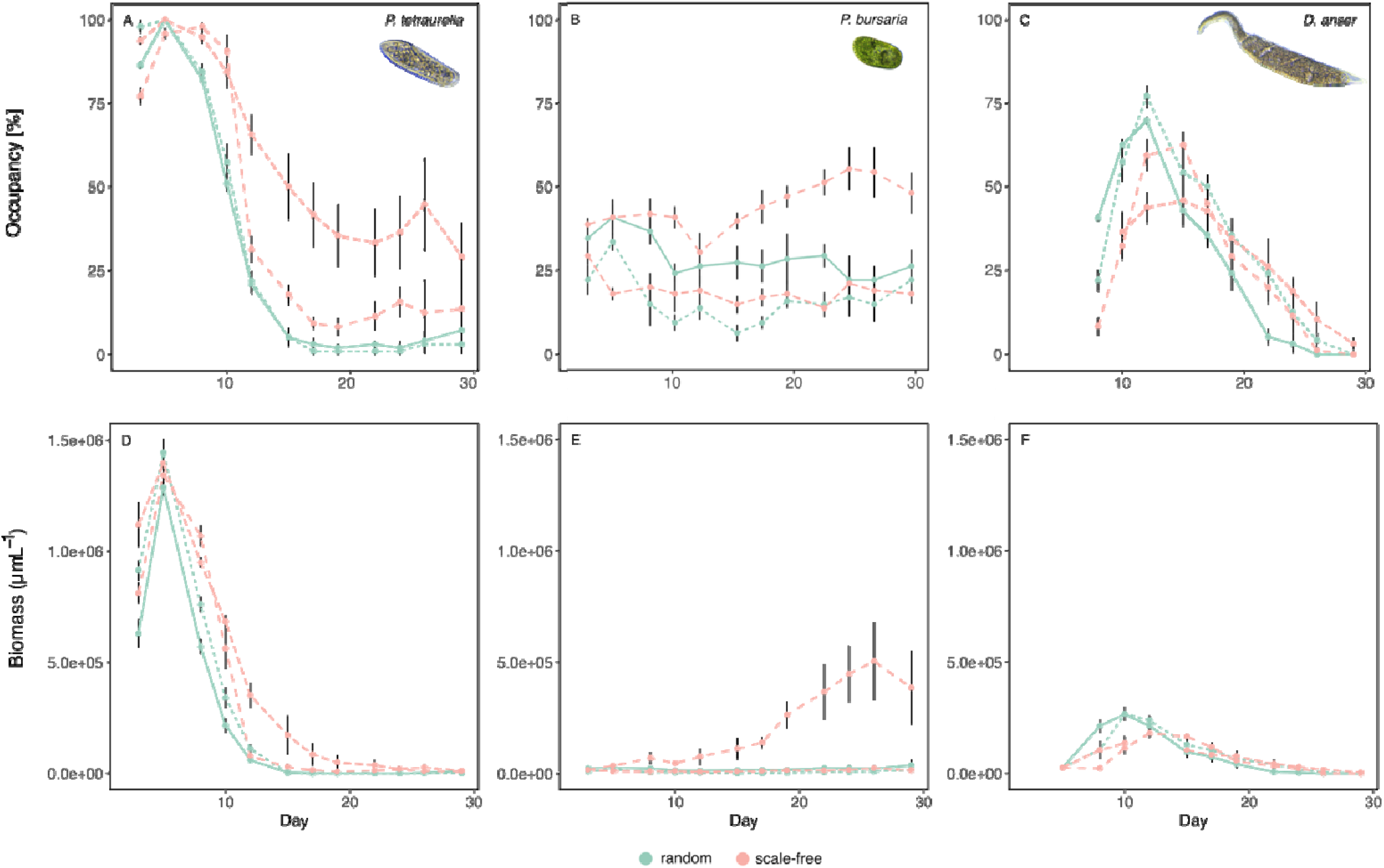
Top panels: Average occupancy (% of occupied patches) over time for (A) *paramecium tetraurelia*, (B) *P. bursaria*, and (C) *Dileptus anser* in experimental metacommunities arranged as scale-free (blue) and random (orange) networks 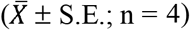. Bottom panels: Average biomass (per metacommunity) over time for (D) *P. tetraurelia*, (E) *P*. . *bursaria*, and (G) *D. anser* in metacommunities arranged as scale-free (light blue) and random (orange) networks 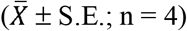. Different lines represent network-level replicates.

### Local dynamics

To address local dynamics within metacommunities, I evaluated the relationship between community descriptors and two topological metrics of patch connectivity.

Metacommunity-level (global) biomass was significantly related to network topology, with both patch metrics showing strong interactions with network type. Biomass declined with increasing degree, and this relationship was stronger for scale-free metacommunities (Fig. 4A-B; GLMM_degree:network_ χ ^2^ = 4.99, *p* << 0.001). This effect is stronger in scale-free networks likely because, architecturally, they present a higher proportion of isolated patches than random networks (Fig. 1). In contrast, closeness centrality had a weaker main effect on biomass but with a similar strong interaction, highlighting a steeper decline in biomass with increasing closeness centrality in scale-free networks (Fig. 4C-D; GLMM_closeness:network_ χ ^2^= 70.867, *p* << 0.001). Overall, degree explained more of the variation in community biomass than closeness centrality (details in supporting information), suggesting that it is a stronger predictor of community-level responses.

**Figure 4.**
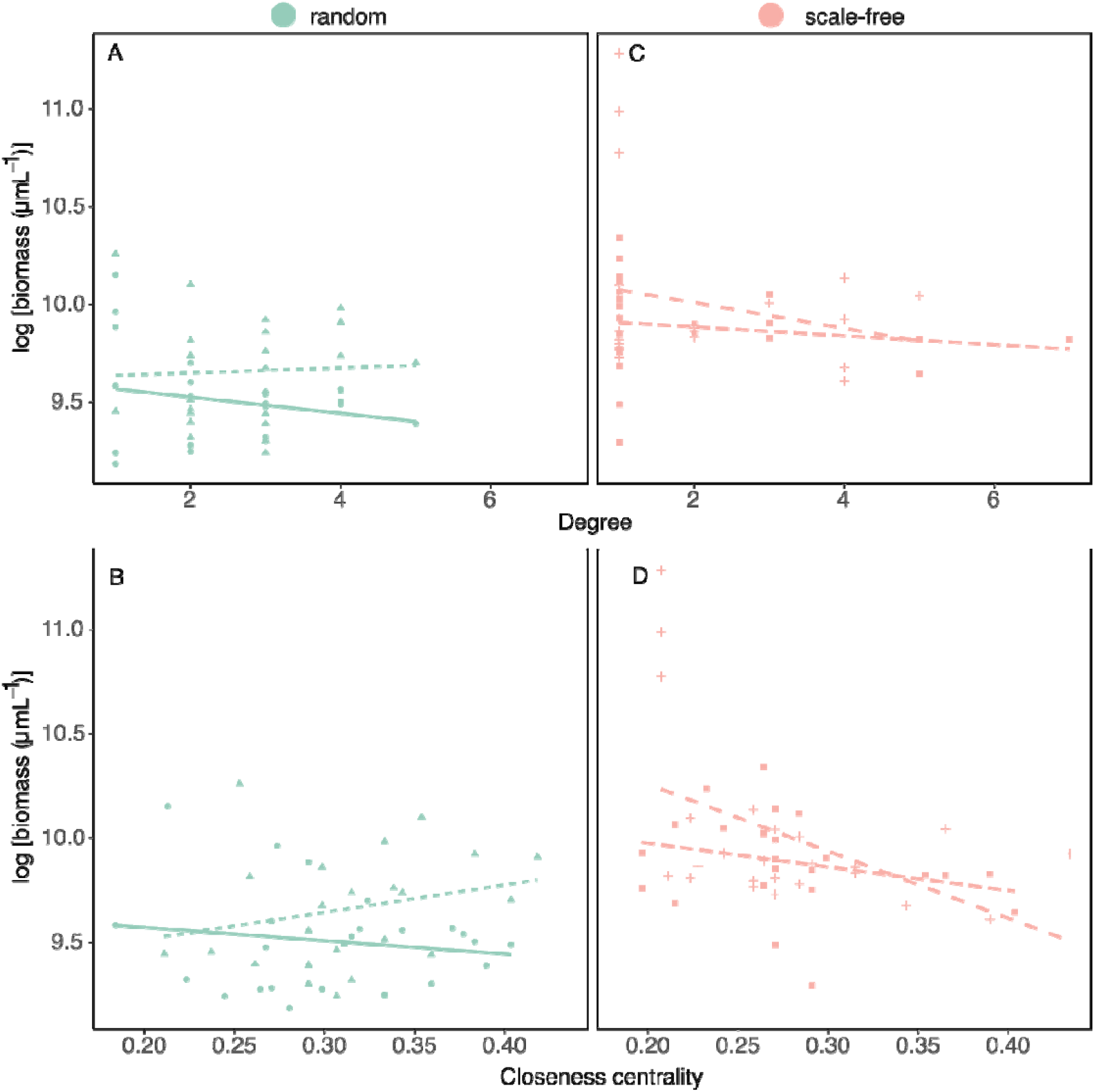
Relationship between local community biomass and (A) degree (i.e., number of neighbors) and (B) patch closeness centrality for metacommunities arranged as random (left ls) and panels scale-free networks (right panels). Different shapes represent different replicate networks.

Patch probability of extinction differed strongly between network types, but showed a contrasting relationship with the different patch metrics. Degree had no effect on the extinction probability in either network type. However, closeness centrality showed a significant interaction with network type. Specifically, patch probability of extinction increased with closeness centrality in random networks but was equal or decreased in scale-free networks (Fig. 5C-D; GLMM_closeness:network_ χ ^2^ = 70.867, *p* << 0.001). These results suggest that more central patches may act as hubs for negative interactions. Specifically, the scale-free topology seems to decouple extinction from spatial position, suggesting that it may provide a structural buffer that stabilizes extinction dynamics regardless of patch centrality.

**Figure 5.**
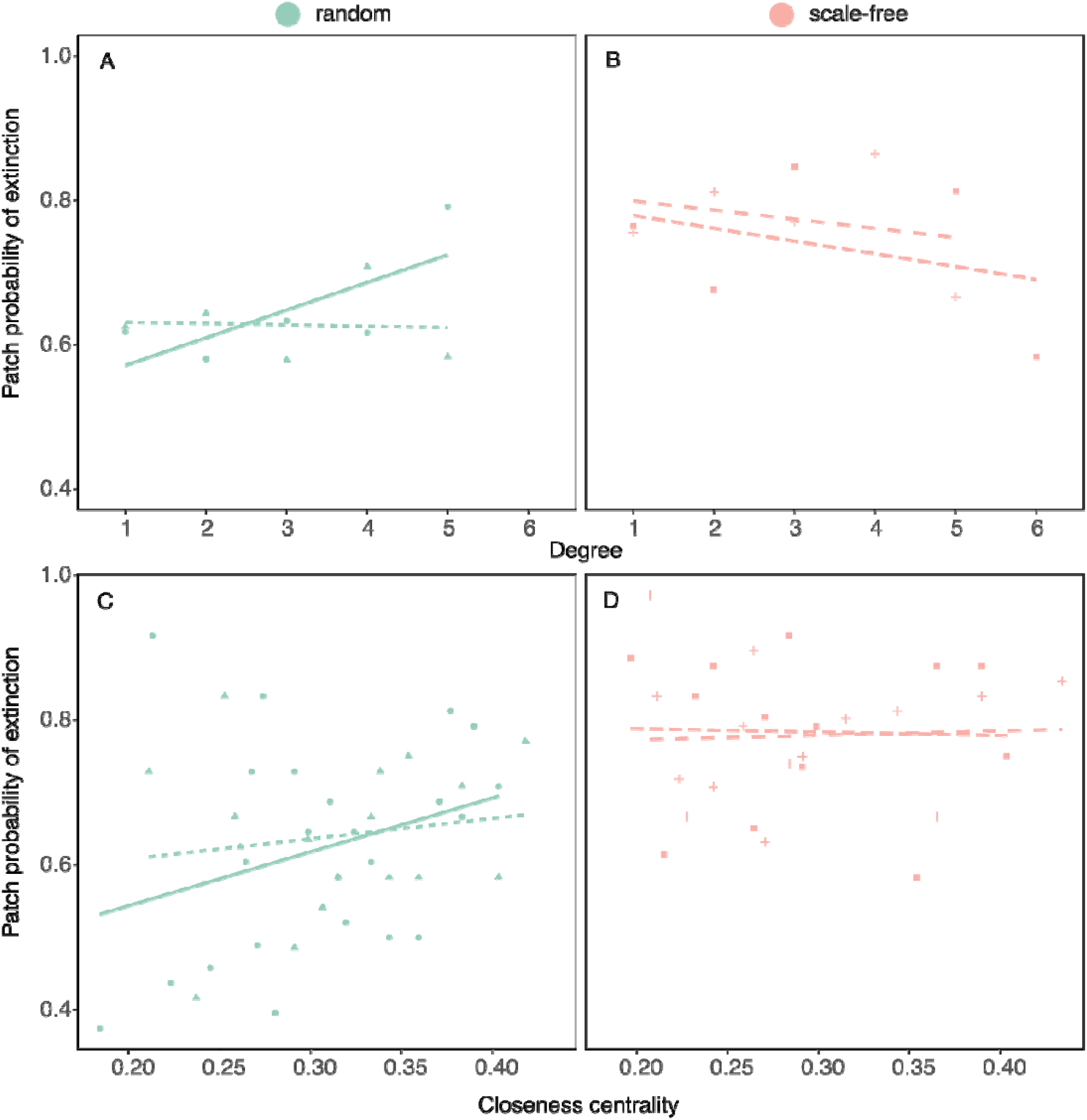
(A) Relationship between probability of extinction of local communities and patch degree (i.e., number of neighbors) in metacommunities arranged as random (left panels) and scale-free networks (right panels). (B) Relationship between probability of extinction of local communities and patch closeness centrality in random (left panels) and scale-free metacommunities (right panels). Each point represents the average among plates (n = 4).

## Discussion

This study shows that network topology does not exert a uniform effect on metacommunities, but instead has scale-dependent effects on ecological processes operating at different spatial scales. While community functioning (i.e. biomass) was primarily associated with local connectivity, extinction dynamics depended on a patch’s position within the network. These results reveal a decoupling among ecological processes, highlighting that different connectivity features drive distinct aspects of metacommunity dynamics.

The community biomass results from this experiment, published in Arancibia (2024), showed that network topology affects community biomass, reaching consistently higher values in scale-free metacommunities. Here, I show that this pattern extends to the species level. The analysis linking community biomass to local patch metrics revealed that community functioning is strongly influenced by local connectivity as captured by patch degree, highlighting the key role of immediate dispersal in regulating local productivity (Leibold *et al*. 2004; Thompson *et al*. 2020). This is also an indication of how metacommunity functioning is sustained by the enhanced functioning of isolated patches, rather than hubs. Because species distributions in metacommunities emerge from the interplay between local interactions and dispersal (Guichard 2017), network topology, by altering the frequency and spatial distribution of interactions should be considered as a key determinant of species distribution.

A comparison with findings in Arancibia & Morin (2022) further illustrates this point. In metapopulations of *P. tetraurelia*, random networks outperformed the scale-free network, likely because increased connectivity enhanced rescue effects. In contrast, all three species in the present metacommunities showed higher biomass in scale-free networks. This suggests that when interactions are present, increased connectivity may instead intensify the negative effects of competition. Scale-free networks with their heterogeneous degree distribution have a higher proportion of low-connectivity patches (Fig. 1). Although this structure weakens rescue effects, it may benefit some species by providing refugia against predators and competitors, allowing them to recover and reach higher biomass, which is consistent with studies that have described a negative relationship between local productivity and dispersal (Mouquet & Loreau 2003). Conversely, the more uniformly high connectivity of random networks, may facilitate the spread of predators or dominant competitors through the network, reducing prey productivity and increasing extinction risk (Borthagaray *et al*. 2015a). In this context patch degree likely captures the primary pathways through which individuals (or resources) move, therefore explaining its relevance for processes operating at local scales.

In contrast to biomass, the patch probability of extinction was not related to local connectivity but instead depended on the patch’s position within the network. This pattern was particularly strong in scale-free metacommunities. These results suggest that persistence is determined not purely by the probability of dispersal but likely by the indirect accessibility of the patch, indicating that being connected to a well-connected patch is more important than the absolute number of local connections. This suggests that extinction dynamics emerge from processes that operate at larger scales than those driving local community functioning.

Altogether, these results reveal that processes within the same metacommunity can respond to different components of network structure. While both community functioning and extinction risk are influenced by dispersal, they respond to different components of network structure, indicating that they operate at different spatial scales. Similar process-dependent effects have been reported by Suzuki & Economo (2024) whose models show that network topology can have variable effects across metacommunity responses, with weaker effects on stability but stronger influences on population size and biodiversity patterns. Together, these findings highlight the need to use multiple connectivity measures to predict different aspects of metacommunity dynamics. Occupancy dynamics results further illustrate the complexity of the effects of network topology, revealing contrasting species-specific responses to network structure. Interspecific differences in the response of community members to spatial connectivity have been documented and can be attributed to differences in key life history traits (Borthagaray *et al*. 2014; Frisch *et al*. 2012; Jones *et al*. 2015). In this case, *P. tetraurelia* consistently showed higher dispersal and reproductive capacity (Arancibia unpublished data). These results are consistent with theoretical expectations that metacommunity dynamics emerge from the interactions between spatial processes and species-specific traits, and highlight the fact that network structure does not act uniformly across all community members. Instead, it can generate heterogeneous responses that can contribute to variation in community dynamics.

Community stability measures in this system showed that local community stability decreases with increasing closeness centrality in scale-free networks, but it is not affected by degree (Arancibia 2024). Altogether, these findings evidence a strong relationship between processes closely linked to persistence, like extinction risk and stability, to network position rather than local connectivity.

These findings have important implications for understanding the effects of habitat fragmentation. Changes in landscape structure and habitat loss not only alter the amount of habitat and its connectivity but also its configuration, which can differentially affect ecological processes such as productivity and persistence. As a result, predicting community responses to environmental change will require multiple measures of connectivity in a spatially explicit context.

## Acknowledgements

I am grateful to Peter Morin, Marcel Holyoak, Julie Lockwood, Nerea Abrego, Cara Faillace, Rita Grunberg, Morin lab members (Rutgers University), and Predictive Community Ecology group members (University of Jyväskylä) for their feedback on different versions of this manuscript.

## References

Amarasekare, P. (2003). Competitive coexistence in spatially structured environments: a synthesis. Ecology Letters, 6, 1109–1122.

Arancibia, P.A. (2024). The topology of spatial networks affects stability in experimental metacommunities. Proceedings of The Royal Society B, 20240567.

Arancibia, P.A. & Morin, P.J. (2022). Network topology and patch connectivity affect dynamics in experimental and model metapopulations. Journal of Animal Ecology, 91, 496–505.

Borthagaray, A.I., Arim, M. & Marquet, P.A. (2014). Inferring species roles in metacommunity structure from species co-occurrence networks. Proceedings of The Royal Society B, 281, 20141425-.

Borthagaray, A.I., Berazategui, M. & Arim, M. (2015a). Disentangling the effects of local and regional processes on biodiversity patterns through taxon-contingent metacommunity network analysis. Oikos, 124, 1383–1390.

Borthagaray, A.I., Pinelli, V., Berazategui, M., Rodríguez-Tricot, L. & Arim, M. (2015b). Effects of metacommunity networks on local community structures: From theoretical predictions to empirical evaluations. Aquatic Functional Biodiversity: An Ecological and Evolutionary Perspective, 75–111.

Brown, B.L., Sokol, E.R., Skelton, J. & Tornwall, B. (2017). Making sense of metacommunities: dispelling the mythology of a metacommunity typology. Oecologia, 183.

Cadotte, M.W. & Fukami, T. (2005). Dispersal, spatial scale, and species diversity in a hierarchically structured experimental landscape. Ecology Letters, 8, 548– 557.

Chisholm, C., Lindo, Z., Gonzalez, A., Chisholm, C., Lindo, Z. & Gonzalez, A. (2011). Metacommunity diversity depends on connectivity and patch arrangement in heterogeneous habitat networks. Ecography, 34, 415–424.

Desjardins, C.D. & Bulut, O. (2015). Journal of Statistical Software Profile Analysis of Multivariate Data in R: An Introduction to the profileR Package. Journal of Statistical Software, 10, 1–29.

Driscoll, D.A. & Lindenmayer, D.B. (2009). Empirical tests of metacommunity theory using an isolation gradient. Ecological Monographs, 79, 485–501.

Economo, E.P. & Keitt, T.H. (2007). Species diversity in neutral metacommunities: a network approach. Ecology Letters, 0, 071117033013001-???

Estrada, E. & Bodin, Ö. (2008). Using network centrality measures to manage landscape connectivity. Ecological Applications, 18, 1810–1825.

Frisch, D., Cottenie, K., Badosa, A. & Green, A.J. (2012). Strong spatial influence on colonization rates in a pioneer zooplankton metacommunity. PLoS ONE, 7, 1– 10.

Gilarranz, L.J. & Bascompte, J. (2012). Spatial network structure and metapopulation persistence. Journal of Theoretical Biology, 297, 11–16.

Gilbert, F., Gonzalez, A. & Evans-Freke, I. (1998). Corridors maintain species richness in the fragmented landscapes of a microecosystem. Proceedings of the Royal Society B: Biological Sciences, 265, 577–582.

Gonzalez, A., Lawton, J.H., Gilbert, F.S., Blackburn, T.M. & Evans-Freke, I. (1998). Metapopulation dynamics, abundance, and distribution in a microecosystem. Science, 281, 2045–2047.

Gonzalez, A., Rayfield, B. & Lindo, Z. (2011). The disentangled bank: How loss of habitat fragments and disassembles ecological networks. American Journal of Botany, 98, 503–516.

Grainger, T.N. & Gilbert, B. (2016). Dispersal and diversity in experimental metacommunities: linking theory and practice. Oikos, 125, 1213–1223.

Guichard, F. (2017). Recent advances in metacommunities and meta-ecosystem theories. F1000Research, 6.

Guzman, L.M., Germain, R.M., Forbes, C., Straus, S., O’Connor, M.I., Gravel, D., et al. (2019). Towards a multi-trophic extension of metacommunity ecology. Ecology Letters, 22, 19–33.

Haddad, N.M., Brudvig, L.A., Clobert, J., Davies, K.F., Gonzalez, A., Holt, R.D., et al. (2015). Habitat fragmentation and its lasting impact on Earth’s ecosystems. Science Advances, 1, e1500052.

Hanski, I. & Gilpin, M. (1991). Metapopulation dynamics: brief history and conceptual domain. Biological Journal of the Linnean Society, 42, 3–16.

Holland, M.D. & Hastings, A. (2008). Strong effect of dispersal network structure on ecological dynamics. Nature, 456, 792–795.

Holt, R.D. (2002). Food web in space: on the interplay of dynamic instability and spatial processes. Ecological Research, 17, 261–273.

Holyoak, M. & Lawler, S.P. (1996). Persistence of an Extinction-Prone Predator-Prey Interaction Through Metapopulation Dynamics. Ecology, 77, 1867–1879.

Holyoak, M., Leibold, M.A. & Holt, R.D. (2005). Metacommunities: Spatial Dynamics and Ecological Communities. The University of Chicago Press, Chicago and London.

Jones, N.T., Germain, R.M., Grainger, T.N., Hall, A.M., Baldwin, L. & Gilbert, B. (2015). Dispersal mode mediates the effect of patch size and patch connectivity on metacommunity diversity. Journal of Ecology, 103.

Kerr, B., Neuhauser, C., Bohannan, B.J.M. & Dean, A.M. (2006). Local migration promotes competitive restraint in a host-pathogen “tragedy of the commons.” Nature, 442, 75–78.

Leibold, M.A., Holyoak, M., Mouquet, N., Amarasekare, P., Chase, J.M., Hoopes, M.F., et al. (2004). The metacommunity concept: A framework for multi-scale community ecology. Ecology Letters, 7, 601–613.

Logue, J.B., Mouquet, N., Peter, H. & Hillebrand, H. (2011). Empirical approaches to metacommunities: A review and comparison with theory. Trends in Ecology and Evolution, 26, 482–491.

MacArthur, R.H. & Wilson, E.O. (1967). The Theory of Island Biogeography. Princeton University Press.

Massol, F., Altermatt, F., Gounand, I., Gravel, D., Leibold, M.A. & Mouquet, N. (2017). How life-history traits affect ecosystem properties: effects of dispersal in meta-ecosystems. Oikos, 126.

Moritz, C., Meynard, C.N., Devictor, V., Guizien, K., Labrune, C., Guarini, J.M., et al. (2013). Disentangling the role of connectivity, environmental filtering, and spatial structure on metacommunity dynamics. Oikos, 122, 1401–1410.

Mouquet, N. & Loreau, M. (2003). Community patterns in source-sink metacommunities. The American naturalist, 162, 544–557.

Santangelo, J.M., Bram Vanschoenwinkel & Trekels, H. (2021). Habitat isolation and the cues of three remote predators differentially modulate prey colonization dynamics in pond landscapes. Oecologia, doi.org/10.1007/s00442-021-04997-6.

Schneider, C.A., Rasband, W.S. & Eliceiri, K.W. (2012). NIH Image to ImageJ: 25 years of image analysis. Nature Methods, 9, 671–675.

Sullivan, L.L., Michalska-Smith, M.J., Sperry, K.P., Moeller, D.A. & Shaw, A.K. (2020). Consequences of ignoring dispersal variation in network models for landscape connectivity. Conservation Biology, 35, 944–954.

Suzuki, Y. & Economo, E.P. (2024). The Stability of Competitive Metacommunities Is Insensitive to Dispersal Connectivity in a Fluctuating Environment. The American Naturalist, 203, 668–680.

Tews, J., Brose, U., Grimm, V., Tielbörger, K., Wichmann, M.C., Schwager, M., et al. (2004). Animal species diversity driven by habitat heterogeneity/diversity: the importance of keystone structures: Animal species diversity driven by habitat heterogeneity. Journal of Biogeography, 31, 79–92.

Thompson, P.L., Guzman, L.M., De Meester, L., Zś, Z., Horváth, Z., Ptacnik, R., et al. (2020). A process-based metacommunity framework linking local and regional scale community ecology.

Tilman, D., Mayt, R.M., Lehman, C.L. & Nowakt, M.A. (1994). Habitat destruction and the extinction debt. Nature, 371, 65–66.

Vanschoenwinkel, B., De Vries, C., Seaman, M. & Brendonck, L. (2007). The role of metacommunity processes in shaping invertebrate rock pool communities along a dispersal gradient. Oikos, 116, 1255–1266.

Warren, P.H. (1996). The effects of between-habitat dispersal rate on protist communities and metacommunities in microcosms at two spatial scales. Oecologia, 132–140.

