## Supplementary materials for "Network topology differentially shapes ecological processes across scales in experimental metacommunities"

### 1. Profile analysis results for occupancy over time

**Community occupancy**

| Test | Statistic | | F | df | p-value |
|---|---|---|---|---|---|
| Parallelism | Hotelling-Lawley | 14.215 | | 11, 4 | 0.06324 |
| Equal levels | - | | 16.95 | 1, 14 | 0.00105 |
| Flatness | - | | 42.13 | 11, 4 | 0.00128 |

***Paramecium tetraurelia* occupancy**

| Test | Statistic | | F | df | p-value |
|---|---|---|---|---|---|
| Parallelism | Hotelling-Lawley | 21.14 | | 11, 4 | 0.03180 |
| Equal levels | - | | 15.21 | 1, 14 | 0.00160 |
| Flatness | - | | 277.21 | 11, 4 | <0.00100 |

***Paramecium bursaria* occupancy**

| Test | Statistic | | F | df | p-value |
|---|---|---|---|---|---|
| Parallelism | Hotelling-Lawley | 3.98 | | 11, 4 | 0.38597 |
| Equal levels | - | | 2.39 | 1, 14 | 0.14400 |
| Flatness | - | | 1.901 | 11, 4 | 0.28097 |

***Dileptus anser* occupancy**

| Test | Statistic | | F | df | p-value |
|---|---|---|---|---|---|
| Parallelism | Hotelling-Lawley | 7.63 | | 8, 6 | 0.02372 |
| Equal levels | - | | 1.25 | 1, 14 | 0.28200 |
| Flatness | - | | - | 8, 7 | - |

### 2. Profile analysis results for biomass over time

***Paramecium tetraurelia* biomass**

| Test | Statistic | | F | df | p-value |
|---|---|---|---|---|---|
| Parallelism | Hotelling-Lawley | 35.98 | | 11, 4 | 0.01204 |
| Equal levels | - | | 17.59 | 1, 14 | 0.00090 |
| Flatness | - | | 196.07 | 11, 4 | <0.00100 |

***Paramecium bursaria* biomass**

| Test | Statistic | | F | df | p-value |
|---|---|---|---|---|---|
| Parallelism | Hotelling-Lawley | 7.17 | | 11, 4 | 0.18374 |
| Equal levels | - | | 4.75 | 1, 14 | 0.04700 |
| Flatness | - | | 4.57 | 11, 4 | 0.07765 |

***Dileptus anser* biomass**

| Test | Statistic | | F | df | p-value |
|---|---|---|---|---|---|
| Parallelism | Hotelling-Lawley | 14.11 | | 8, 6 | 0.00498 |
| Equal levels | - | | 5.69 | 1, 14 | 0.03180 |
| Flatness | - | | - | 8, 7 | - |

3. **Model definition and output for the relationship between community biomass and degree (lmer package)**

Initially, this model included random intercepts for network identity (net_name) and plate, but since they did not explain any variance, the final model retained only day as a random effect.

community_biomass ~ degree*network_type + (1|day)

**Model summary:**

N=960; Day=12; Network_name = 4; Plate = 4

$R^2$ (marginal)= 0.189; $R^2$ (conditional)= 0.48

| Fixed effects | Estimate | SE | t value | $\chi^2$ | p($\chi^2$) |
|---|---|---|---|---|---|
| Intercept | 121374 | 27495 | 4.414 | | |
| Degree | -16392 | 3630 | -4.516 | 270.421 | <<0.0001 |
| Network (scale-free) | 145070 | 15982 | 9.077 | 41.306 | <<0.0001 |
| Degree x network | -31572 | 4572 | -6.906 | 47.688 | <<0.0001 |

| Random effects | Variance | SD |
|---|---|---|
| Day | $7.333e^{+09}$ | 85630 |
| Residual | $1.265e^{+10}$ | 112466 |

4. **Model definition and output for the relationship between community biomass and closeness centrality**

community_biomass ~ closeness_centrality*network_type + (1|day) + (1|net_name) +(1|plate)

**Model summary:**

N=3264; Day=12; Network_name = 4; Plate = 4

$R^2$ (marginal)= 0.075; $R^2$ (conditional)= 0.333

| Fixed effects | Estimate | SE | t value | $\chi^2$ | p($\chi^2$) |
|---|---|---|---|---|---|
| Intercept | 23889 | 9394 | 2.543 | | |
| Closeness centrality | -16929 | 16061 | -1.054 | 88.069 | <<0.0001 |
| Network (scale-free) | 76866 | 8331 | 9.226 | 15.659 | <<0.0001 |
| Degree x network | -197107 | 23414 | -8.418 | 70.867 | <<0.0001 |

| Random effects | Variance | SD |
|---|---|---|
| Day | $6.419e^{+08}$ | 25335 |
| Network name | $1.785e^{+07}$ | 4224 |
| Plate | $1.624e^{+05}$ | 403 |
| Residual | $1.660e^{+09}$ | 40744 |

5. **Model definition and output for the relationship between patch probability of extinction (PE) and patch degree.**

Initially, this model included random intercepts for network identity (net_name), but since they did not explain any variance, the final model retained only plate as a random effect.

PE ~ closeness_centrality*network_type + (1|plate)

**Model summary:**

N=384; Plate = 4

$R^2$ (marginal)= 0.148; $R^2$ (conditional)= 0.186

| **Fixed effects** | **Estimate** | **SE** | **t value** | $\chi^2$ | $p(\chi^2)$ |
|---|---|---|---|---|---|
| Intercept | 0.592752 | 0.039189 | 15.126 | | |
| Closeness centrality | 0.010635 | 0.012101 | 0.879 | 0.6161 | 0.4325 |
| Network (scale-free) | 0.173787 | 0.040143 | 4.329 | 66.8228 | <<0.0001 |
| Degree x network | -0.007715 | 0.015054 | -0.512 | 0.2626 | 0.6083 |

| **Random effects** | **Variance** | **SD** |
|---|---|---|
| Plate | 0.001555 | 0.03943 |
| Residual | 0.032606 | 0.18057 |

6. **Model definition and output for the relationship between patch probability of extinction (PE) and closeness centrality.**

Initially, this model included random intercepts for network identity (net_name), but since they did not explain any variance, the final model retained only plate as a random effect.

PE ~ closeness_centrality*network_type + (1|plate)

**Model summary:**

N=384; Plate = 4

$R^2$ (marginal)= 0.160; $R^2$ (conditional)= 0.198

| **Fixed effects** | **Estimate** | **SE** | **t value** | $\chi^2$ | $p(\chi^2)$ |
|---|---|---|---|---|---|
| Intercept | 0.43543 | 0.07734 | 5.630 | | |
| Closeness centrality | 0.60048 | 0.23932 | 2.509 | 2.0690 | 0.15032 |
| Network (scale-free) | 0.36373 | 0.09926 | 3.664 | 70.3031 | <<0.0001 |
| Degree x network | -0.69802 | 0.33257 | -2.099 | 4.4052 | 0.0358 |

| **Random effects** | **Variance** | **SD** |
|---|---|---|
| Plate | 0.00156 | 0.03949 |
| Residual | 0.03213 | 0.17925 |

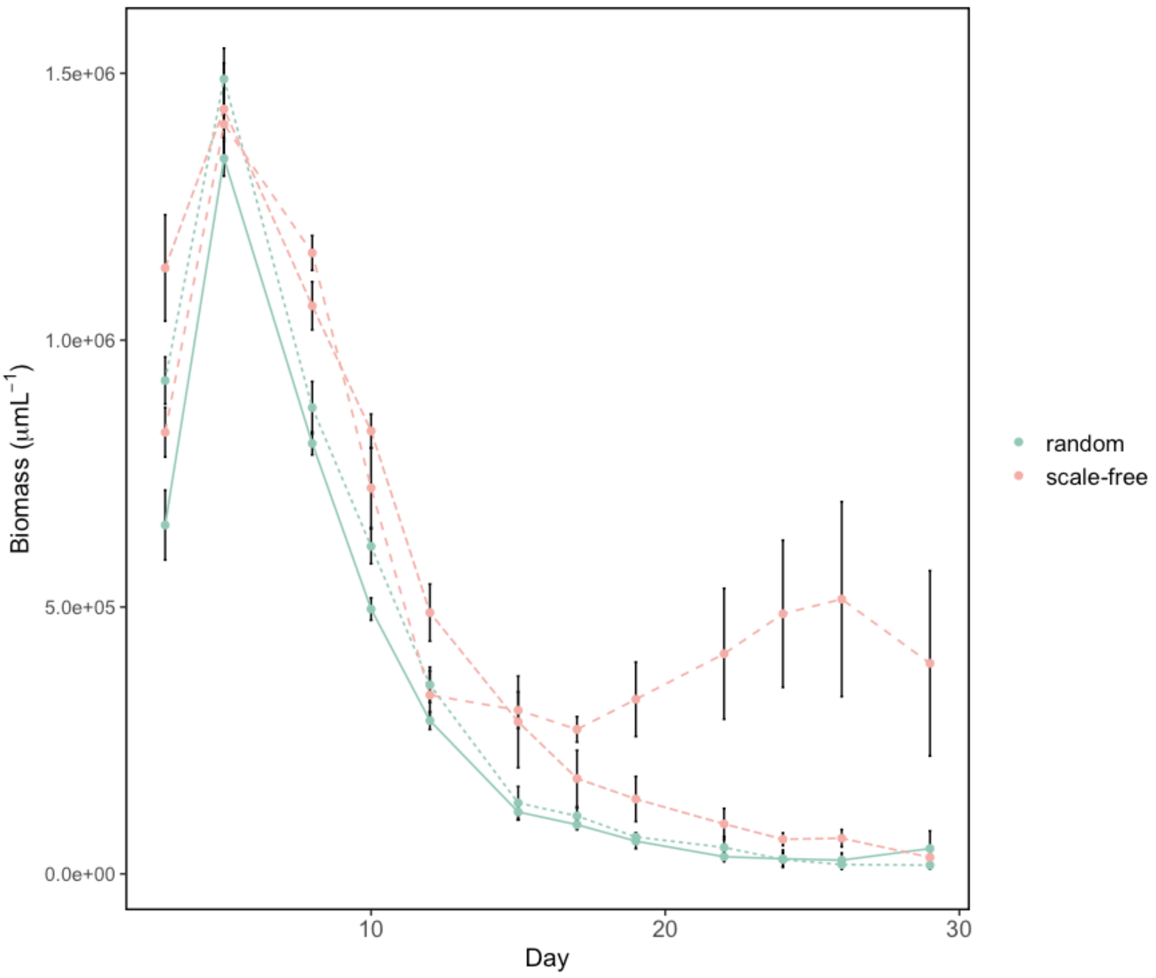


**Fig. S1**. Metacommunity biomass. Redrawn using data from Arancibia (2024). Values are identical to those presented in the original publication.
